# A Deep Learning Approach for Histology-Based Nuclei Segmentation and Tumor Microenvironment Characterization

**DOI:** 10.1101/2022.12.08.519641

**Authors:** Ruichen Rong, Hudanyun Sheng, Kevin W. Jin, Fangjiang Wu, Danni Luo, Zhuoyu Wen, Chen Tang, Donghan M. Yang, Liwei Jia, Mohamed Amgad, Lee A.D. Cooper, Yang Xie, Xiaowei Zhan, Shidan Wang, Guanghua Xiao

## Abstract

Microscopic examination of pathology slides is essential to disease diagnosis and biomedical research; however, traditional manual examination of tissue slides is laborious and subjective. Tumor whole-slide image (WSI) scanning is becoming part of routine clinical procedure and produces massive data that capture tumor histological details at high resolution. Furthermore, the rapid development of deep learning algorithms has significantly increased the efficiency and accuracy of pathology image analysis. In light of this progress, digital pathology is fast becoming a powerful tool to assist pathologists.

Studying tumor tissue and its surrounding microenvironment provides critical insight into tumor initiation, progression, metastasis, and potential therapeutic targets. Nuclei segmentation and classification are critical to pathology image analysis, especially in characterizing and quantifying the tumor microenvironment (TME). Computational algorithms have been developed for nuclei segmentation and TME quantification within image patches; however, existing algorithms are computationally intensive and time-consuming for WSI analysis.

In this study, we present Histology-based Detection using Yolo (HD-Yolo), a new method that significantly accelerates nuclei segmentation and TME quantification. We demonstrate that HD-Yolo outperforms existing methods for WSI analysis in nuclei detection and classification accuracy, as well as computation time.

## 1. Introduction

Histopathology examination of tissues is the cornerstone of disease diagnosis and prognosis. In this procedure, a pathologist examines cells, their spatial organizations and tissue structures under a microscope seeking for hints of disease and its effect on the tissue. It requires experienced pathologists to identify and interpret subtle morphological patterns in histopathology slides. Manual examination of pathology slides is laborious and time consuming; moreover, results are subjective among hospitals and individuals. With advances in imaging technology, whole-slide image (WSI) scanning of tissue slides is becoming a routine clinical procedure. Advanced whole-slide scanners are capable of rapidly producing massive numbers of WSIs that capture histopathological details in high resolution. As a result, digital pathology has seen increasing adoption in clinical practice and diagnosis. Recently, deep learning algorithms have developed and used for pathology image analysis^1–3^. The advanced algorithms have been applied in primary diagnosis^4–6^, prognosis studies^7,8^, as well as association analyses between pathology image and genomic data^9^.

A tumor is a mass of tissue with complex structures consisting of cancer cells and surrounding non-malignant cells, which form the tumor micro-environment (TME). Study and characterize TME can provide critical insights into tumor initiation, progression, metastasis, and potential therapeutic targets. Nuclei segmentation and classification is a key step in digital pathology image analysis. Several algorithms have been developed to precisely identify nuclei in pathology image analysis, which enable researchers to characterize and quantify TME by extracting TME-related imaging features and associating these features with patient outcomes and genomic information. For example, Wang et al. proposed <u>H</u>istology-based <u>D</u>igital (HD)-Staining algorithm for nuclei localization, classification, and masking. HD-Staining relies on the Mask R-CNN^10^ instance segmentation algorithm. Mask R-CNN is constructed from the following components: first, a deep CNN-based backbone with a feature pyramid network (FPN)^11^ to extract imaging features from the raw image; second, a region proposal network (RPN)^12^ to localize potential objects; third, a detection header to classify detected objects into different types; and fourth, a mask header to segment the boundary of detected objects. Since Mask R-CNN processes object localization and classification separately, it is considered a two-stage object detection algorithm; alternatively, algorithms that merge the second and third components are considered one-stage object detection algorithms. In earlier years, one-stage object detection algorithms had the advantage in computational efficiency, while two-stage object detection algorithms achieved better accuracy and coverage by sacrificing speed, especially on small objects. Using Mask R-CNN as its main nuclei segmentation algorithm, HD-Staining takes hours to analyze a single slide and is therefore inefficient at analyzing large-scale datasets.

To reduce the computational cost of nuclei segmentation, Graham et al. proposed HoVer-Net^13^, an approach that directly applied three distinct branches atop image features extracted by a VGG-style backbone: a nuclear classification (NC) branch, a nuclear pixel (NP) branch, and a HoVer branch. HoVer-Net relied on energy-based post-processing to generate nuclei masks from the outputs of the NP branch and HoVer branch. Due to its network design and memory cost, HoVer-Net accepts relatively small patches (80×80) as input and takes significant computational time for its post-processing masks.

Both HD-Staining and HoVer-Net algorithms perform well at analysis of pathology image patches. However, their computational requirements are prohibitively large for WSIs, which has greatly limited their application to WSI analysis. Recent developments in deep learning have brought forth more efficient object detection algorithms applicable to varying speed and accuracy requirements. For example, Cascade R-CNN^14^ provides high-quality two-stage detection for dense and occluded objects; EfficientDet^15^ improves detection efficiency and accuracy under resource constraints by automatically selecting network designs through a compound scaling approach, and achieved the highest performance in comparison to other networks across multiple benchmarks by publication time; FCOS^16^ simplifies one-stage detection by completely avoiding anchor box operations; DETR^17^ is an end-to-end object detector by directly applying a transformer encoder-decoder on top of feature maps. Accounting for algorithm stability and efficiency, the most widely used one-stage detection algorithms are that of the Yolo family^18^. With advanced training and network design, (Scaled-)Yolov4^19,20^, YoloR^21^, YoloX^22^, as well as Yolov6^23^, and Yolov7^24^ enable one-stage detection algorithms to achieve comparable performance to that of two-stage algorithms.

In this study, we developed Histology-based Detection Yolo (HD-Yolo), an algorithm for nuclei detection and classification as well as TME-related feature extraction for WSI analysis. HD-Yolo enhances computational efficiency and enriches slide-level TME features. The HD-Yolo system consists of a nuclei classification and segmentation algorithm and an efficient TME feature extraction method designed for WSI analysis. HD-Yolo provides three key contributions: Firstly, HD-Yolo enables simultaneous detection and segmentation by including a segmentation module within the traditional Yolo architecture. Secondly, HD-Yolo significantly increases the computation speed and reduces the requirement for computational resources WSI analysis. Thirdly, HD-Yolo utilizes a density-based TME feature extraction module to summarize WSI-level and ROI-level features. We demonstrated the performance of HD-Yolo by applying it to different tissue types: lung cancer, liver tissue, and breast cancer. We further applied HD-Yolo to the Cancer Genome Atlas - Breast Cancer (TCGA-BRCA) dataset^25^ and illustrated the prognostic value of image features extracted by HD-Yolo through survival analysis. Finally, to demonstrate its usage, we developed a web frontend that implements HD-Yolo and displays WSI results.

## 2. Materials and Methods

### 2.1 Datasets

#### 2.1.1 Lung adenocarcinoma dataset

The lung adenocarcinoma dataset curated in the HD-Staining paper^26^ contains 127 patches (500×500) extracted from 39 pathological ROIs in the Lung Screening Trial (NLST) dataset^27^. A board-certified pathologist (L.J.) manually annotated nuclei type and masks in these image patches. Image patches were split into training, validation, and testing sets based on slide ID. Specifically, 105 patches from 29 slides were assigned to training, 12 patches from another 5 slides are assigned to validation, and the other 10 patches from the remaining 5 slides were reserved for testing. Nuclei were manually labeled and segmented by pathologists into 6 different categories: tumor nuclei, stromal nuclei, lymphocyte nuclei, macrophage nuclei, red blood cells, and karyorrhexis. More than 12,000 cell nuclei were included in the training set (tumor nuclei 24.1%, stromal nuclei 23.9%, lymphocytes 29.5%, red blood cells 5.8%, and others 16.7%), while 1227 and 1086 nuclei were included in the validation and testing sets, respectively^26^.

#### 2.1.2 Liver tissue dataset

The liver tissue dataset contains 76 hepatic H&E slides (40× magnification) in liver tissues. This dataset consisted of 2 normal mouse liver slides, 9 normal human liver slides, 2 cirrhotic human liver slides without hepatocellular carcinoma (HCC), and 63 cirrhotic human liver slides with HCC. We selected 51 image patches (500×500) in non-malignant regions and manually labeled nuclei into six categories: hepatocyte nuclei, stroma nuclei, lymphocyte nuclei, and red blood cells. The 51 image patches were split into 35 for training, 8 for validation, and 8 for testing, respectively.

#### 2.1.3 Breast cancer dataset

The publicly available NuCLS dataset^28^ consists of many image patches extracted from breast cancer images from The Cancer Genome Atlas (TCGA) dataset^25^. Image patches were split into single-rater and multi-rater datasets. Nuclei within the patches were annotated through the collaborative effort of pathologists, pathology residents, and medical students. The corrected single-rater dataset was collected with high-quality annotations supervised by one pathologist and further split into 5 different folds (fold1-fold5) for training and validating nuclei detection and segmentation algorithms. The multi-rater dataset is a relatively smaller cohort validated by multiple pathologists and is therefore, suitable as an independent testing dataset. The corrected single-rater dataset contains 1744 image patches, 54916 annotations with class labels, and 27976 unlabeled annotations, while the multi-rater evaluation dataset contains 53 image patches, 1203 annotations with class labels, and 150 unlabeled annotations. The nuclei were assigned into 20 subcategories and four super-classes: tumor, stromal, stromal tumor-infiltrating lymphocytes (sTILs), and others. All models compared in this study were trained and validated on fold1 in the corrected single-rater dataset and further evaluated on the multi-rater dataset. Here we report super-class performances for ease of comparison with other models, as well as the statistics reported in the original papers.

#### 2.1.4 TCGA – BRCA dataset

We analyzed the prognosis of breast carcinoma based on the H&E slides and patient clinical information available in the TCGA-BRCA dataset. Among all 1,061 patients with pathology images, 653 IDC (Infiltrating Ductal Carcinoma) and 169 ILC (Infiltrating Lobular Carcinoma) female patients with confirmed tumor stage information were used in this study. Patients were split into 80% training (658 patients) and 20% testing (164 patients) stratified by histological types. **Supplementary Table 1** summarizes the patient demographic and clinical information in the training and testing datasets.

### 2.2 Methodology in HD-Yolo framework

Figure 1. illustrates the overall workflow of the HD-Yolo system. For each slide, we first divided the whole tissue region into small patches (640×640 at 40× magnification). We used a Yolo-based object segmentation algorithm (**Figure 1a**) to quickly locate nuclei coordinates, identify nuclei type, and extract shape morphological features. We then summarized the above information into spatial TME features with the feature extraction pipeline (**Figure 1b**). The feature extraction pipeline focuses on selected image patches in the ROI and analyzes the entire slide for a comprehensive characterization of the TME.

**Figure 1.**
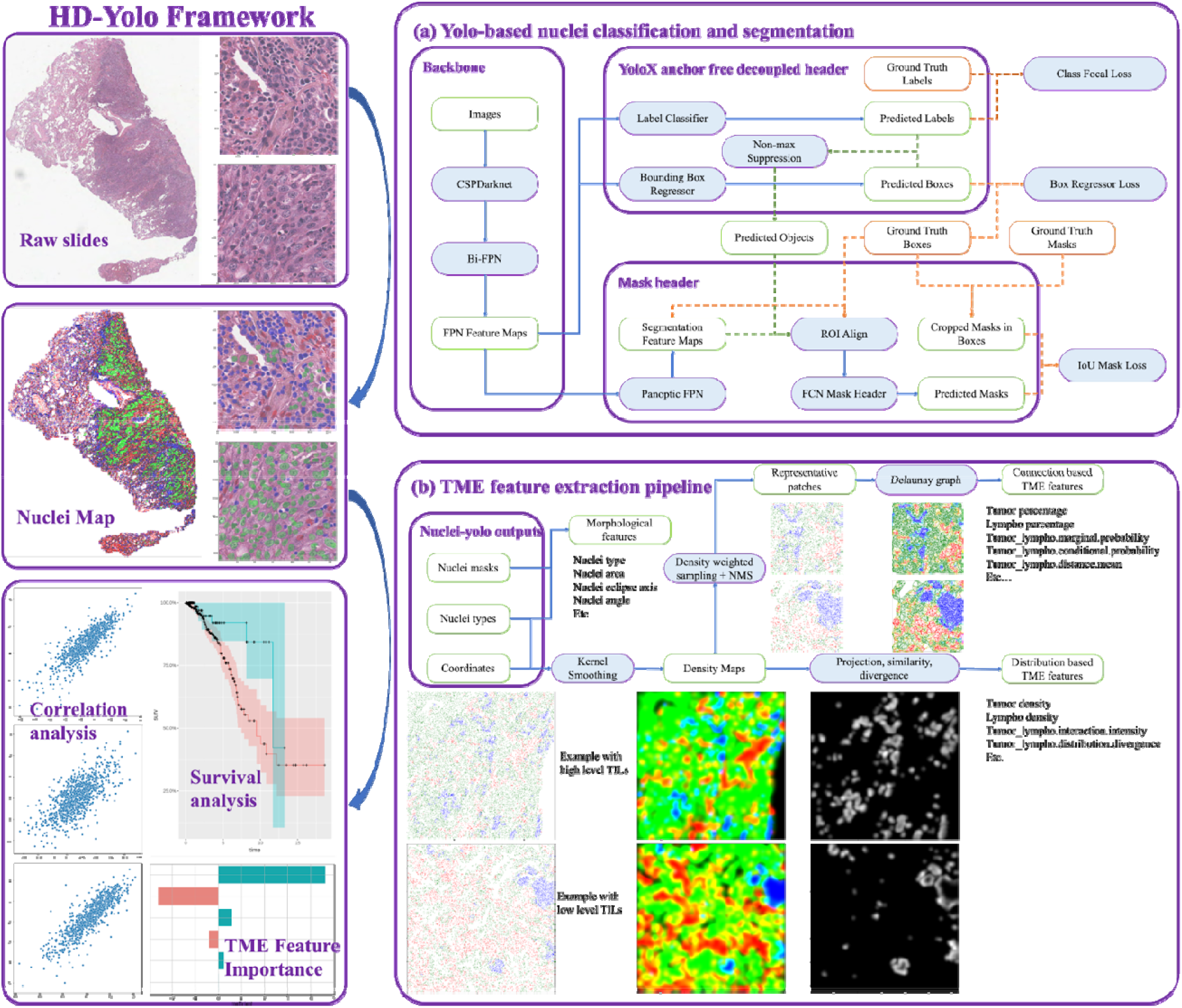
HD-Yolo system workflow. (a) displays the nuclei detection and segmentation model in the HD-Yolo system. The model is constructed from three components: a Bi-FPN backbone, a YoloX detection header, and an FCN mask header. The light blue boxes with purple borders are deep learning modules. The white boxes with green borders are intermediate outputs of the model. The solid blue lines are computational graphs used in both training and inference. The orange dotted lines are only processed in training while the green dotted lines are for inference only. (b) illustrates the TME feature extraction pipeline based on the nuclei density maps built from the nuclei detection results. The pipeline directly extracts whole-slide level TME features (distributions, projections, similarity, and divergence) from the spatial densities of different types of nuclei. The pipeline also summarizes ROI-level TME features by automatically selecting ROIs from the above analysis.

#### 2.2.1 HD-Yolo nuclei segmentation model architecture

The nuclei detection and segmentation model in the HD-Yolo system was constructed under the Yolo architecture with a customized design as illustrated in **Figure 1a**. We used Yolov5’s CSPDarkNet as the backbone for image feature extraction^20^. CSPDarkNet employs Cross Stage Partial Network (CSPNet) layers to partition and merge the original feature maps in DarkNet for cross-stage communication. We then applied the BiFPN^15^ network to blend backbone feature maps at different resolutions. We further utilized the YoloX decoupled detection head^22^ to localize and classify objects. As one-stage object detection algorithms, the Yolo family cannot directly provide object masks for detected objects; therefore, we utilized the Panoptic FPN^29^ to build feature maps for segmentation and added a one-class fully convolutional network (FCN) head to output masks for detected nuclei. For our task, the YoloX decoupled detection head achieved better accuracy than the merged head used in Yolov3-Yolov5, and the decoupled head was capable of further splitting nuclei location and type information for representative analysis. The segmentation head provides rich information for both nuclei masking and background segmentation and was faster than the multi-RoIAlign + multi-class CNN header used in Mask R-CNN. Note that with the mask header added, HD-Yolo is not strictly considered a one-stage object detection algorithm, but its overall computational speed is competitive with that of the latest Yolo variations without mask support and is still much faster than existing nuclei segmentation algorithms.

#### 2.2.2 WSI nuclei density-based TME feature extraction

Due to computational speed limitations, most existing TME analysis systems are deployed on a limited number of randomly selected patches when analyzing large-scale high-resolution slides. For instance, the HD-Staining pipeline characterizes nuclei spatial organization by building Delaunay triangle graphs in 100 patches (1024×1024) randomly sampled from the ROI. This approach relies on human-labeled ROIs and the results would be potentially biased according to the size and number of available patches inside the ROI. When ROIs are large, important subregions may be ignored from calculation; on the other hand, when ROIs are small, overlapped subregions are likely to be analyzed multiple times. To alleviate the overlapping issue, one may sample a small number (10, for instance) of larger patches (2048×2048) to include more connections in a single image, but this increases variance during summarization; on the other hand, sampling a higher number (100, for instance) of smaller patches (1024×1024) reduces between-patch variance but excludes connections at the patch border from the calculation. Our HD-Yolo system enables fast nuclei segmentation over whole slides and thus makes summarizing spatial properties globally feasible. We refined the procedure of Delaunay triangulation by automatically selecting ROI based on nuclei densities without human labeling. In addition to ROI-based TME features, we expanded several slide-level TME features by analyzing nuclei distribution and densities over all nuclei in the slide (**Figure 1b**).

The following comprises HD-Yolo’s TME feature extraction process. First, the nuclei segmentation algorithm is run on whole slides and the coordinates, types, and sizes of all detected nuclei are recorded. Each type of nuclei is then allocated into a 2D point cloud and kernel smoothing^30^ is applied based on nuclei size to generate a density map. These density maps reflect the distribution of different types of nuclei in the slide. To extract slide-level TME features, the following intensity features are calculated for each pair of nuclei *type_i* and *type_j*, based on their density maps *map_i* and *map_j*: the dot-product between *map_i* and *map_j*, which summarizes the overall interactions between *type_i* and *type_j*; the projection from *map_i* to *map_ j*, which represents the level of *type_j* influenced by *type_i* and vice versa; and the cosine similarity between *map_i* and *map_j*, which represents the interaction intensity between *type_i* and *type_j*. These density features are irrelevant to the size and number of patches sampled and are therefore robust to ROI size variation. To extract ROI-based TME features, the above analysis is utilized to automatically decide ROIs and build a Delaunay triangle graph in each region. By default, regions with high tumor densities and regions with high interaction intensities are potential ROIs. Delaunay triangle triplets are calculated in each region, and the edge probabilities for each pair of nuclei *type_i* and *type_j* are averaged over all regions.

### 2.3 HD-Yolo performance evaluation criteria

#### 2.3.1 Evaluation of nuclei detection and segmentation performance

We compared HD-Yolo with existing models using the following statistics: 1) Detection coverage: the percentage of ground truth nuclei detected by the model; 2) Nuclei classification accuracy: the percentage of ground truth nuclei that were further correctly classified by the model; 3) Matthews correlation coefficient (MCC): a score of the model’s overall performance, considering accuracy and coverage among different classes; 4) Median Intersection-over-Union (mIoU): the mIoU between the predicted mask and the original mask measures the segmentation similarity between detected nuclei and ground truth nuclei; 5) Precision: the percentage of true nuclei among detected nuclei; 6) Recall rate: how many ground return nuclei are detected; 7) F1-score: the harmonic mean of precision and recall; 8) Time per image: the mean inference speed for the whole dataset on a Tesla V100 GPU with 32 GB of memory. Coverage, accuracy, MCC, precision, recall, and F1-score were evaluated at thresholding, with the IoU of the ground truth box and prediction box larger than 0.5. For precision, recall, and F1-score, we eliminated the unbalancing between each class by calculating the class-specific statistics and then averaging the scores over all classes. For time per image, we recorded the total computational time from when the image was loaded into the model to when the results of the whole dataset were exported, and then averaged over the number of images. More specifically, we included the post-processing time of each model in the calculation, while file reading, plotting, and result exporting times were excluded. We selected the highest batch size and number of processors to allow maximal computational resources (GPU, CPU, etc.) for each model.

#### 2.3.2 Evaluation of WSI analysis and feature extraction

We compared the WSI inference and feature extraction times of different algorithms when processing slides of different sizes from the BRCA dataset. The regularized Cox proportional hazard model^31–33^ was used to analyze prognosis hazard ratios and the ElasticNet algorithm was used for feature selection. We applied two prognosis analyses separately on: 1) IDC patients only, and 2) all IDC and ILC patients. As there were zero death events for ILC in the testing dataset, we did not perform separate analyses for ILC. For each dataset, we built multiple survival models with different sets of features: 1) Clinical features only; 2) Clinical features plus Delaunay graph-based TME features defined in HD-Staining; 3) Clinical features plus density-based TME features as defined in Section 2.2.2; and 4) ElasticNet-selected features from all features listed above.

For each survival model, we first utilized a grid-search strategy to select the best hyperparameters according to the 10-fold cross-validation C-index and then applied the best model to the testing dataset. We report the cross-validation C-index in the training dataset, the training C-index with standard error, and the testing C-index with standard error to measure the performance of each survival model. The feature coefficients, permutation importance, and p-values are also provided to demonstrate feature selection results.

#### 2.4 HD-Yolo implementation details

HD-Yolo was developed in Python 3.9. The Yolo-based nuclei segmentation algorithm was modified from the official YoloX (https://github.com/MegEngine/YOLOX) and Yolov5 repositories (https://github.com/ultralytics/yolov5, commit: 19c8760caa70f4d04d3fb974d797f9d922bf6eb8). We modified the Bi-FPN backbone and YoloX header with our mask support and immigrated the model into the Yolov5 distributed data infrastructure. We further reimplemented the training, validation, and testing pipeline to support intensive data augmentation, joint detection, and segmentation training, and customized data with missing and unclassified labels/masks, as well as whole slide inference. Major packages used in HD-Yolo development and evaluation include PyTorch 1.9.0, torchvision 0.10.0, scikit-image 0.18.1, scikit-learn 0.24.1, and openslide 3.4.1. For training and inference, we distributed the data across 4 × 32 GB Tesla V100 GPUs and deployed the model on a Tesla V100 GPU. The regularized CoxPH model was built with the scikit-survival 0.17.2 package and the permutation importance score was calculated with ELI5 0.11.0 in Python. The survival statistics C-index and p-value were calculated with the R packages survival, glmnet, and survcomp in R 3.6.1.

For comparison purposes, we implemented the following algorithms as described in their original publications and default settings in GitHub: 1) HD-Staining (Mask R-CNN with resnet50 backbone, https://github.com/matterport/Mask_RCNN, commit: 3deaec5d902d16e1daf56b62d5971d428dc920bc); 2) Mask R-CNN: we customized the PyTorch Mask R-CNN with EfficientNet-B3 backbone^34^ and softNMS^35^ for better accuracy and faster speed; 3) HoVer-Net (https://github.com/vqdang/hover_net); 4) Yolov7:

https://github.com/WongKinYiu/yolov7, 5) Yolov6: https://github.com/meituan/YOLOv6, 6) ScaledYolov4: https://github.com/WongKinYiu/ScaledYOLOv4, 7) FCOS: https://github.com/Adelaide-AI-Group/FCOS; 8) Deformable-DETR: https://github.com/open-mmlab/mmdetection; 9) EfficientDet: https://github.com/rwightman/efficientdet-pytorch; 10) Cascade R-CNN: https://github.com/open-mmlab/mmdetection.

## 3 Results and Discussion

### 3.1 HD-Yolo nuclei segmentation performance

We compared HD-Yolo’s performance on the lung cancer dataset with that of three existing nuclei segmentation algorithms: HD-Staining, PyTorch Mask R-CNN, and HoVer-Net. All models were evaluated using 500×500 image patches at 40× magnification. Nuclei were detected, segmented, and classified into 6 categories: tumor nuclei, stromal nuclei, lymphocyte nuclei, macrophage nuclei, red blood cells, and karyorrhexis. The performance and speed of each model on the testing and validation datasets are listed in **Table 1** and **Supplementary Table 2**.

**Table 1.** Comparison of nuclei detection performance and speed among different models on lung cancer testing dataset. HD-Staining was processed with the repository and hyperparameters from the original paper. HoVer-Net was finetuned with the pre-trained model from the PanNuke dataset^36^ released by the author. Mask R-CNN was upgraded from the official TorchVision Mask R-CNN by replacing the ResNet-50 FPN backbone with an EfficientNet-B3 FPN backbone for greater accuracy and faster speed. HD-Yolo achieved the fastest inference with the best performance.

| Testing | Coverage | Accuracy | Precision | Recall | F1-score | mIoU | Time per image (s) |
| --- | --- | --- | --- | --- | --- | --- | --- |
| HD-Yolo | **0.8263** | **0.7110** | **0.8308** | **0.6743** | **0.7409** | **0.8423** | **0.0108** |
| HD-Staining | 0.7724 | 0.6553 | 0.7943 | 0.6310 | 0.6994 | 0.8390 | 0.1503 |
| Mask R-CNN | 0.8000 | 0.6864 | 0.8215 | 0.6454 | 0.7198 | 0.8419 | 0.1704 |
| HoVer-Net | 0.7851 | 0.5867 | 0.6599 | 0.3953 | 0.5808 | 0.7717 | 1.0400 |

We observed that HD-Yolo achieved superior performance and speed among all models: HD-Yolo has ∼5% greater coverage on ground truth nuclei than HD-Staining and HoVer-Net; HD-Yolo achieves ∼6% higher accuracy than HD-Staining and ∼12% higher accuracy than HoVer-Net; and HD-Yolo is 15 times faster than HD-Staining (0.01s vs. 0.15s) and 100 times faster than HoVer-Net (0.01s vs. 1s).

For the task of nuclei segmentation, HD-Yolo generated smooth, round masks for different types of nuclei and provided accurate masks for occluded nuclei. HD-Yolo’s mIoU is competitive with that of HD-Staining and Mask R-CNN and is significantly higher than that of HoVer-Net. As shown in in **Figure 2**, we observed that HoVer-Net failed to segment occluded nuclei and tended to split one large nucleus into several smaller ones. We ascribe the success of our segmentation header to the following reasons: firstly, detection features are generally distinct from segmentation features, especially for one-stage detection algorithms, and Panoptic FPN provides important intrinsic image features for precise segmentation; secondly, nuclei morphological features are relatively similar across different types, and a one-class shallow fully-convolutional network (FCN) network can provide satisfactory masks at considerable low computational cost; thirdly, FCN-based mask headers (utilized by HD-Yolo, HD-Staining, and Mask R-CNN) are more robust than heuristic post-processing (HoVer-Net) on irregularly-shaped nuclei and occluded nuclei.

**Figure 2.**
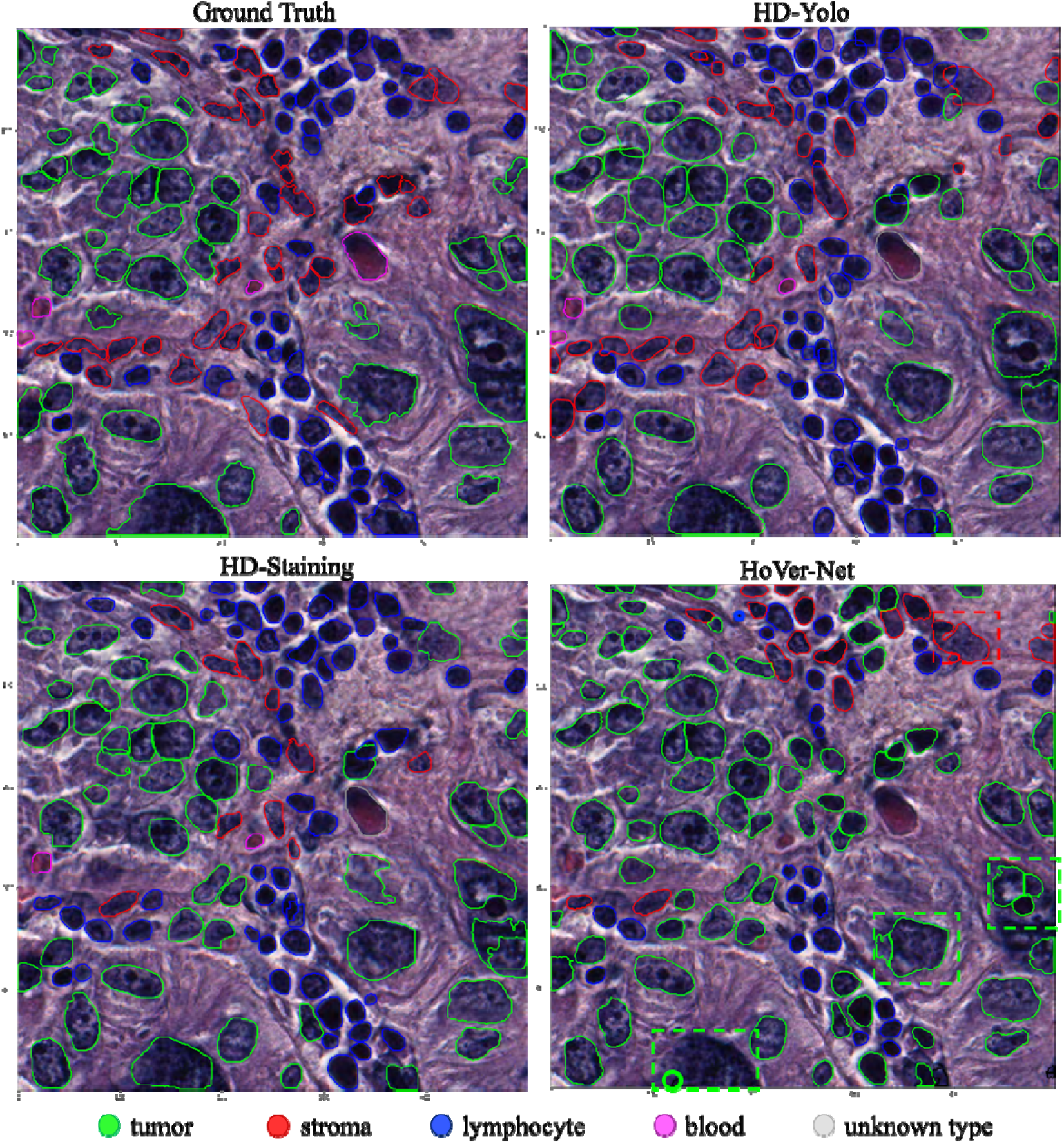
Nuclei segmentation results for different models on a testing lung cancer image patch. HD-Yolo and HD-Staining generated round and smooth nuclei-shaped masks. HoVer-Net failed to segment occluded objects, a very large tumor nucleus (bottom left rectangle), and tended to split large nuclei into smaller pieces (examples are annotated with dashed bounding boxes).

We further employed transfer learning to build a liver nuclei detection model from the above lung cancer HD-Yolo model (**Figure 3a**). The liver cancer dataset contained only 35 training, 8 validation, and 8 testing patches, which is relatively small compared to the lung cancer study. The model achieved 0.9287 coverage and 0.8437 accuracy on the testing dataset after only 10 epochs of fine-tuning (**Figure 3b**), demonstrating HD-Yolo’s generalizability.

**Figure 3.**
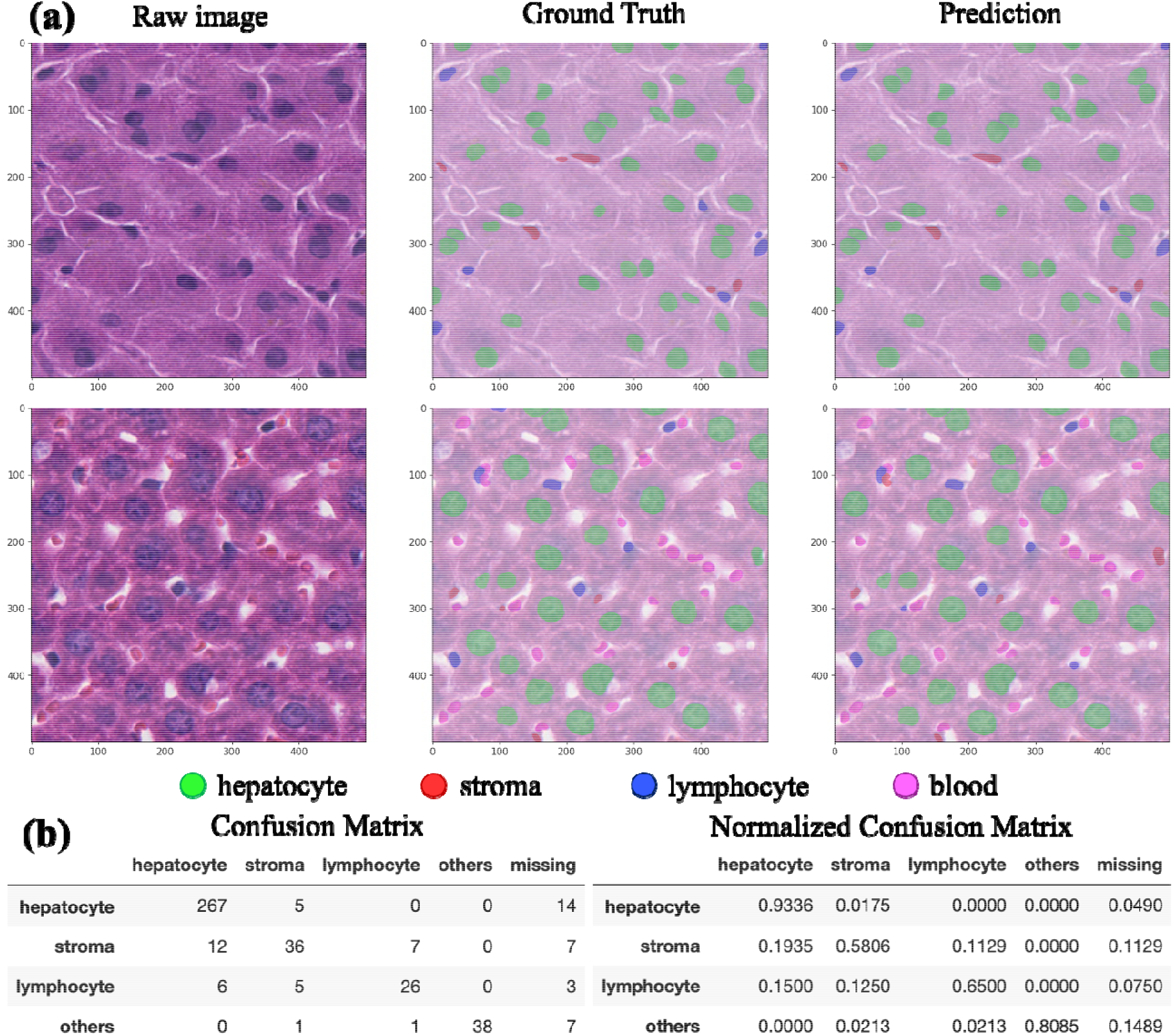
Liver HD-Yolo nuclei detection model transferred from the lung cancer model. (a) Two examples in the testing dataset. (b) Raw and normalized confusion matrix of the testing dataset.

### 3.2 HD-Yolo object detection performance

In addition, we compared HD-Yolo detection performance to that of advanced object detection algorithms on the large-scale NuCLS benchmark dataset. The NuCLS dataset contains missing labels and lacks complete mask annotations; moreover, not all models included in this comparison were able to provide masks without extra modification. We, therefore, focus only on detection performance based on three predefined super-classes: tumor, stromal, and sTILs. As shown in **Table 2**, HD-Yolo achieved similar performance to that of a variety of advanced object detection algorithms. We observed that one-stage algorithms (HD-Yolo, Yolov6, and Yolov7) and two-stage algorithms (EfficientDet and Cascade R-CNN) achieved the highest F1-score; however, HD-Yolo, Yolov6, and Yolov7 outperformed other models in computational speed. Surprisingly, FCOS and deformable-DETR maintained comparable precision, but had significantly lower coverage compared to other methods. This could be due to FCOS and deformable-DETR’s utilization of heatmap and transformer-based approaches, which are not suitable for cases where the same type of nuclei at different sizes are densely overlapped. Overall, we believe that HD-Yolo can achieve state-of-the-art performance compared to currently available object detection algorithms and providing reliable results for breast cancer WSI analysis.

**Table 2.** Nuclei detection performance on the NuCLS multi-rater testing dataset. The HD-Yolo model is competitive with other advanced object detection algorithms, achieving blazing fast inference speed.

| Network | Backbone | Coverage | Accuracy | Precision | Recall | F1-score | Time per image (s) |
| --- | --- | --- | --- | --- | --- | --- | --- |
| HD-Yolo | CSPDarkNet-l-BiFPN | **0.8950** | 0.8042 | 0.9069 | **0.7992** | 0.8493 | 0.0089 |
| Yolov7 | E-ELAN | 0.8723 | 0.8013 | 0.9065 | 0.7945 | 0.8468 | 0.0090 |
| Yolov6 | Efficient-Rep-PAN | 0.8798 | **0.8133** | **0.9101** | 0.7963 | **0.8494** | **0.0088** |
| Scaled-Yolov4 | CSPDarkNet-PANet | 0.8603 | 0.7977 | 0.8764 | 0.7810 | 0.8249 | 0.0123 |
| FCOS | ResNet101-FPN | 0.7876 | 0.7574 | 0.8848 | 0.7556 | 0.8148 | 0.0381 |
| Deformable-DETR | ResNet50 | 0.7889 | 0.7620 | 0.8877 | 0.7588 | 0.8180 | 0.1420 |
| EfficientDet | EfficientNetB3-BiFPN | 0.8726 | 0.7950 | 0.8930 | 0.7936 | 0.8403 | 0.0951 |
| Mask R-CNN | ResNet101-FPN | 0.8631 | 0.7881 | 0.8749 | 0.7828 | 0.8252 | 0.1462 |
| Cascade R-CNN | ResNeXt101_32x4d-FPN | 0.8230 | 0.7915 | 0.9103 | 0.7902 | 0.8460 | 0.1720 |

### 3.3 HD-Yolo WSI analysis

#### 3.3.1 HD-Yolo WSI inference with single and ensembled models

Ensemble modeling is a common method of improving overall inference performance by combining multiple models without altering the network structure. To build a more accurate breast cancer model, we ensembled 5 independent nuclei segmentation models trained and validated on 5 predefined splits (fold1-fold5) in the NuCLS corrected-single rater dataset. During inference, the input images went through 5 different models and duplicated detections were filtered by non-max suppression. We further halved inference speed without influencing performance by switching from single-precision (float32) models to half-precision (float16) models. **Table 3** shows the performance and speed of the ensembled model as well as every independent model on the inferred P-truth testing dataset. The ensemble model achieved even higher performance, with a ∼5% higher detection rate (coverage), ∼7% higher detection accuracy, ∼6% higher recall, and ∼7% higher MCC compared to the single models. Therefore, we believe that both the single models and the ensembled model can be generalized to large-scale breast cancer whole slide analysis for TME feature extraction; single models may be utilized for fast screening, while the ensembled model can provide more accurate nuclei detection results. Performance for each super-class (tumor, stromal, and sTILs) and confusion matrices are summarized in **Supplementary Table 3** and **Supplementary Table 4**.

**Table 3.** Performance of cross-validation study and final ensemble model on the multi-rater inferred P-truth testing dataset. The variance between models trained on different folds is rarely small. Ensemble modeling further improved performance by ∼5% at the expense of computational speed.

|  | Coverage | Accuracy | Precision | Recall | F1-score | MCC | mIoU | Time per image (s) |
| --- | --- | --- | --- | --- | --- | --- | --- | --- |
| Model1 | 0.8914 | 0.8017 | 0.9063 | 0.7960 | 0.8474 | 0.7246 | 0.7550 | 0.0042 |
| Model2 | 0.8891 | 0.7958 | 0.8989 | 0.7840 | 0.8373 | 0.7156 | 0.7545 | 0.0040 |
| Model3 | 0.8965 | 0.8092 | 0.9030 | 0.8026 | 0.8498 | 0.7335 | 0.8019 | 0.0041 |
| Model4 | 0.8995 | 0.8101 | 0.9012 | 0.8023 | 0.8478 | 0.7339 | 0.7725 | 0.0043 |
| Model5 | 0.8987 | 0.8050 | 0.8910 | 0.8050 | 0.8446 | 0.7288 | 0.7946 | 0.0041 |
| Mean | 0.8950 | 0.8044 | 0.9001 | 0.7980 | 0.8454 | 0.7273 | 0.7757 | 0.0041 |
| Ensembled | 0.9364 | 0.8655 | 0.9238 | 0.8655 | 0.8937 | 0.8044 | 0.8191 | 0.0197 |

We analyzed all H&E slides in the TCGA breast cancer dataset and compared inference speeds between HD-Yolo single, HD-Yolo ensemble, HD-Staining, and HoVer-Net. For HD-Yolo and HD-Staining, we roughly extracted the tissue region from each slide by thresholding and scanned the slides with a 512×512 window. For HoVer-Net, we used the default tissue extraction and patch preparation pipeline as defined in the original repository. For HD-Staining and HoVer-Net, we used the maximum possible batch size that fit in a 32 GB memory GPU and the maximum amount of CPU for post-processing. For HD-Yolo, we limited GPU memory to < 8 GB (for a 512×512 window with a batch size of 64) and used at most 64 CPU threads (32 core) to mimic a small server setup with balanced CPU and GPU resources. An example of HD-Yolo’s whole slide inference results can be found in **Supplementary Figure 1. Table 4** summarizes whole slide inference speeds for all models on the BRCA dataset based on tissue sizes under different quantiles: minimum, median, 95% quantile, and maximum. The HD-Yolo single model took only 1/30 to 1/25 of the time used by HD-Staining and 1/130 to 1/50 of the time used by HoVer-Net. The more accurate ensemble HD-Yolo model took double the processing time of the single version. Accounting for the substantial processing time expended when exporting the morphological results of tens of thousands of nuclei, HD-Yolo’s overall speed is about 10 to 15 times faster than HD-Staining and 50 times faster than HoVer-Net. It is noteworthy that in this comparison, HD-Yolo uses only a fourth of the computational resources used by HD-Staining and HoVer-Net but is already 50 to 100 times faster. We expect even greater improvement on large slides with more GPU memory and CPU cores by increasing image patch size, batch size, and I/O workers. As the majority of slides (> 95% quantile) took less than 5 minutes to process with the single model and ∼10 minutes with the ensemble model, the whole dataset (1061 slides) took about 12 hours with the single model and 1 day with the ensemble model on a server with four GPUs. The same task would take 1 to 2 weeks with HD-Staining and longer with HoVer-Net.

**Table 4.** Inference speed comparison between different models on TCGA breast cancer dataset. HD-Yolo single model is about 25 to 30 times faster than HD-Staining and 50 to 130 times faster than HoVer-Net. The HD-Yolo ensemble model took double the time of its single counterpart with slightly better accuracy.

| Quantiles | Minimum | Median | 95% quantile | Maximum |
| --- | --- | --- | --- | --- |
| No. patches (512×512) | 803 | 15891 | 22552 | 51969 |
| ROI diameter (40×, pixel) | 16.38k | 72.85k | 86.78k | 131.72k |
| HD-Yolo ensemble (h:m:s) | 00:00:43 | 00:09:23 | 00:15:31 | 0:26:41 |
| HD-Yolo single (h:m:s) | 00:00:19 | 00:03:39 | 00:06:58 | 0:12:00 |
| HD-Staining (h:m:s) | 00:07:33 | 01:43:08 | 02:24:18 | 5:37:22 |
| HoVer-Net (h:m:s) | 00:15:50 | 04:46:38 | 11:51:22 | 26:38:12 |
| TME feature extraction | 00:00:22 | 00:03:12 | 00:04:47 | 00:06:52 |
| Export tables and figures | 00:01:09 | 00:29:15 | 00:32:56 | 00:45:58 |

#### 3.3.2 TCGA breast cancer survival analysis based on clinical and nuclei spatial features

We extracted TME features based on the nuclei segmentation results from the fast HD-Yolo single model computed in section 3.2.1 and built a CoxPH model for breast cancer prognosis. For each slide, we constructed a spatial density map based on the nuclei type, location, and shape morphological features of nuclei belonging to the three super-classes: tumor, stromal, sTILs. We include in the analysis the 15 Delaunay graph-based interaction features defined in HD-Staining as well as the 12 nuclei density-based nuclei TME features in HD-Yolo defined in section 2.2.2. In addition to the image-based TME features, the following clinical features are included in the analysis: age at diagnosis, AJCC pathologic tumor stage (AJCC tumor stage), estrogen receptor (ER) status by immunohistochemistry (ER status by IHC), progesterone receptor status by immunohistochemical (PR status by IHC), human epidermal growth factor receptor 2 by immunohistochemical (HER2 status by IHC). Detailed information regarding the TME and clinical features can be found in **Supplementary Table 5**.

We applied the regularized CoxPH model to analyze prognosis hazard ratios and used the ElasticNet algorithm for feature selection. The cross-validation C-index, training C-index with standard error, and the testing C-index with standard error for different models and feature sets can be found in **Figure 4a**. We observed that, despite the clinical features of age at diagnosis and tumor stage, TIL-related TME features defined by HD-Yolo are highly correlated with survival outcomes (**Supplementary Table 6** provides feature importance scores and p-values): *l_t*.*proj*.*prob* has the highest negative coefficient while *t_l*.*proj*.*prob* has the second highest positive coefficient other than age (**Figure 4b**). The *l_t*.*proj*.*prob* (p-value < 0.05) feature represents the degree to which tumor tissue has been invaded by sTILs. A high *l_t*.*proj*.*prob* indicates that large numbers of tumors in tissue regions have been invaded by sTILs and are strongly interacting with the lymphocytes. This is consistent with the prognostic model that an increase in *l_t*.*proj*.*prob* will likely reduce the hazard ratio and lead to better survival outcomes. On the other hand, the *t_l*.*proj*.*prob* (p-value < 0.01) feature represents the degree to which sTILs are surrounded by tumor cells. A high *t_l*.*proj*.*prob* indicates that lymphocytes have been overwhelmed by surrounding tumor nuclei and thus cannot effectively inhibit tumor tissue growth. This is consistent with the prognostic model that an increase in *t_l*.*proj*.*prob* will likely increase the hazard ratio and lead to worse survival outcomes.

**Figure 4.**
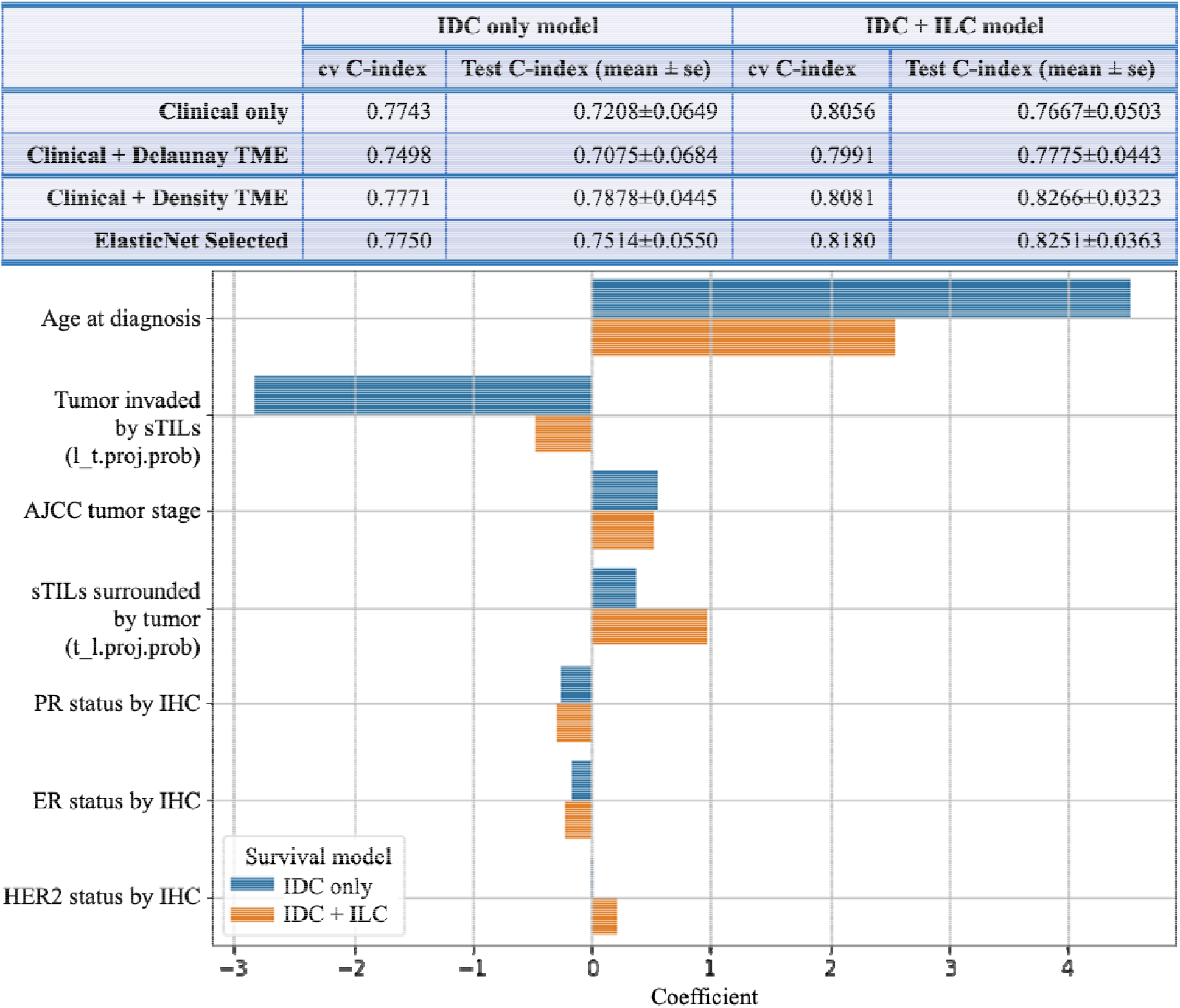
CoxPH survival analysis for breast cancer subtypes. The top panel reports the cross-validation C-index and testing C-index under different datasets and feature sets. Clinical features include age at diagnosis, AJCC pathologic tumor stage (AJCC tumor stage), estrogen receptor (ER) status by immunohistochemistry (ER status by IHC), progesterone receptor (PR) status by immunohistochemical (PR status by IHC), human epidermal growth factor receptor 2 by immunohistochemical (HER2 status by IHC). Clinical + Delaunay TME uses clinical features with Delaunay graph-based features. Clinical + Density TME includes clinical features with nuclei density-based features. ElasticNet Selected takes clinical features plus all image features with ElasticNet permutation importance > 0 and p-value < 0.05. The bottom panel plots the coefficients of the selected features from ElasticNet.

## 4 Conclusion

In this paper, we proposed HD-Yolo, a system for fast, accurate nuclei segmentation and rich TME feature extraction. Our comparison studies show that the HD-Yolo algorithm outperformed existing algorithms (Mask R-CNN, HoVer-Net, etc.) in nuclei detection and segmentation accuracy. HD-Yolo also provided high-quality masks without sacrificing efficiency, in contrast to advanced object detection algorithms. With its fast nuclei detection algorithm, HD-Yolo significantly accelerated WSI analysis while using less computational resources. In addition, the TME features summarized by HD-Yolo’s nuclei density-based method are highly correlated with patient prognosis and provide meaningful explanations regarding tumor progression and survival. The efficient and accurate HD-Yolo pipeline can be potentially useful for a variety of tasks in pathology imaging analysis. In summary, we expect HD-Yolo to be a powerful tool that will facilitate the analysis of digital pathology and provide meaningful biological insights.

## Data availability

For lung cancer analysis, the pathology patches were extracted from the NLST dataset (https://cdas.cancer.gov/nlst/). For breast cancer, pathology patches and annotations that were used for algorithm training, validation, and comparison are available in the online NuCLS portal (https://sites.google.com/view/nucls/). The whole slide imaging support the findings of survival study is available at The Cancer Genome Atlas - Breast Cancer (TCGA-BRCA, https://www.cancer.gov/types/breast).

## Acknowledgements

This work has been supported by the National Institutes of Health [R01GM140012, R01GM141519, R01DE030656, U01CA249245, R35GM136375, P50CA070907, 2P30CA142543], and the Cancer Prevention and Research Institute of Texas [CPRIT RP180805 and RP190107]. The funding bodies had no role in the design, collection, analysis, or interpretation of data in this study.

