## Supplementary Figures and Tables for "A Deep Learning Approach for Histology-Based Nuclei Segmentation and Tumor Microenvironment Characterization"

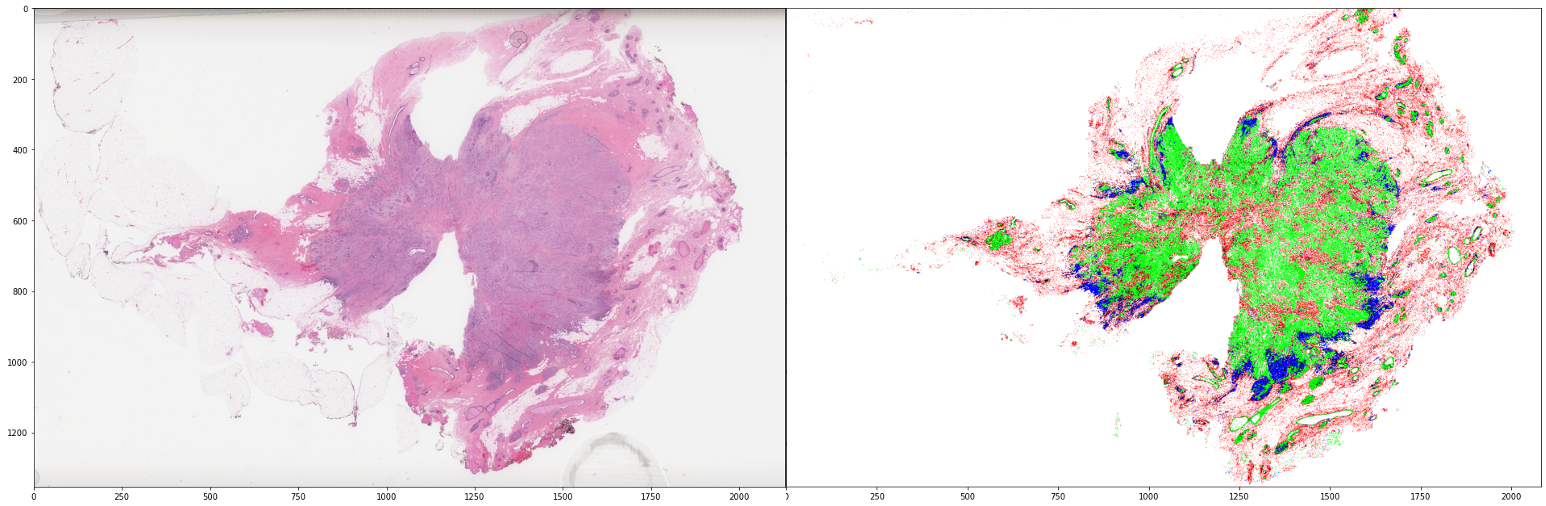

Supplementary Figure 1. Results of HD-Yolo inference on an example breast cancer (BRCA) slide. The whole slide and inference results have been shrunk to 1/16 of their original size for display purposes. Green: tumor nuclei; red: stromal nuclei; blue: sTILs; pink: red blood cells.

| Demographics | Type | Train (658) | Test (164) | G-test p-value |
| --- | --- | --- | --- | --- |
| Histological Type | IDC | 523 | 130 | 0.9625 |
|  | ILC | 135 | 34 |  |
| Stage | Stage I | 114 | 30 | 0.1521 |
|  | Stage II | 387 | 86 |  |
|  | Stage III | 149 | 42 |  |
|  | Stage IV | 8 | 6 |  |
| Age | <40 | 43 | 15 | 0.2400 |
|  | 40-55 | 233 | 50 |  |
|  | 55-70 | 254 | 73 |  |
|  | 70+ | 128 | 26 |  |
| Race | White | 476 | 121 | 0.6807 |
|  | Black | 89 | 22 |  |
|  | Others | 34 | 5 |  |
|  | Unknown | 59 | 16 |  |
| Ethnicity | Hispanic | 23 | 8 | 0.6353 |
|  | Non-Hispanic | 524 | 126 |  |
|  | Unknown | 111 | 30 |  |

Supplementary Table 1. Demographic and clinical information of the training and testing dataset for survival analysis. P-values were calculated with G-test across the training and testing datasets.

| Testing | Coverage | Accuracy | Precision | Recall | F1-score | mIoU | Time per image (s) |
| --- | --- | --- | --- | --- | --- | --- | --- |
| HD-Yolo | 0.7985 | 0.6387 | 0.8180 | 0.6270 | 0.7033 | 0.8359 | 0.0164 |
| HD-Staining | 0.7757 | 0.6085 | 0.7981 | 0.6191 | 0.6913 | 0.8330 | 0.1535 |
| Mask R-CNN | 0.8026 | 0.6079 | 0.7715 | 0.5836 | 0.6495 | 0.8274 | 0.1855 |
| HoVer-Net | 0.7167 | 0.4380 | 0.6212 | 0.3931 | 0.4130 | 0.7007 | 1.0500 |

Supplementary Table 2. Nuclei detection performance on the lung cancer validation dataset for different models.

|  | Precision | | | Recall | | | F1-score | | | MCC | | |
| --- | --- | --- | --- | --- | --- | --- | --- | --- | --- | --- | --- | --- |
|  | tumor | stromal | sTILs | tumor | stromal | sTILs | tumor | stromal | sTILs | tumor | stromal | sTILs |
| Model1 | 0.9287 | 0.8385 | 0.9482 | 0.8274 | 0.7722 | 0.7919 | 0.8751 | 0.8040 | 0.8630 | 0.7959 | 0.7381 | 0.8154 |
| Model2 | 0.9289 | 0.8358 | 0.9346 | 0.8552 | 0.7089 | 0.8108 | 0.8905 | 0.7671 | 0.8683 | 0.8176 | 0.6962 | 0.8189 |
| Model3 | 0.9507 | 0.8322 | 0.9301 | 0.8413 | 0.7532 | 0.8270 | 0.8926 | 0.7907 | 0.8755 | 0.8260 | 0.7217 | 0.8269 |
| Model4 | 0.9474 | 0.8550 | 0.8932 | 0.8214 | 0.7089 | 0.8811 | 0.8799 | 0.7751 | 0.8871 | 0.8076 | 0.7092 | 0.8367 |
| Model5 | 0.9659 | 0.8047 | 0.9093 | 0.7877 | 0.7563 | 0.8676 | 0.8678 | 0.7798 | 0.8880 | 0.7973 | 0.7041 | 0.8396 |
| Mean | 0.9443 | 0.8332 | 0.9231 | 0.8266 | 0.7399 | 0.8357 | 0.8812 | 0.7833 | 0.8764 | 0.8089 | 0.7138 | 0.8275 |
| Ensemble | 0.9512 | 0.8814 | 0.9226 | 0.8889 | 0.8228 | 0.8703 | 0.9190 | 0.8511 | 0.8957 | 0.8643 | 0.8006 | 0.8513 |

Supplementary Table 3. Per-class performance of single models under different train/val splits, as well as the ensemble models. We provide the precision, recall, F1-score, and MCC for each super-class: tumor, stromal, and sTILs.

| **Ensemble** | | | | | **Model1** | | | | |
| --- | --- | --- | --- | --- | --- | --- | --- | --- | --- |
|  | **tumor** | **stromal** | **sTILs** | **missing** |  | **tumor** | **stromal** | **sTILs** | **missing** |
| **tumor** | 448 | 16 | 2 | 38 | **tumor** | 417 | 13 | 2 | 72 |
| **stromal** | 16 | 260 | 25 | 15 | **stromal** | 27 | 244 | 14 | 31 |
| **sTILs** | 7 | 19 | 322 | 22 | **sTILs** | 5 | 34 | 293 | 38 |
| **Model2** | | | | | **Model3** | | | | |
|  | **tumor** | **stromal** | **sTILs** | **missing** |  | **tumor** | **stromal** | **sTILs** | **missing** |
| **tumor** | 431 | 16 | 2 | 57 | **tumor** | 424 | 19 | 5 | 56 |
| **stromal** | 25 | 224 | 21 | 46 | **stromal** | 16 | 238 | 18 | 44 |
| **sTILs** | 8 | 28 | 300 | 34 | **sTILs** | 6 | 29 | 306 | 29 |
| **Model4** | | | | | **Model5** | | | | |
|  | **tumor** | **stromal** | **sTILs** | **missing** |  | **tumor** | **stromal** | **sTILs** | **missing** |
| **tumor** | 414 | 26 | 4 | 60 | **tumor** | 397 | 42 | 5 | 60 |
| **stromal** | 20 | 224 | 35 | 37 | **stromal** | 11 | 239 | 27 | 39 |
| **sTILs** | 3 | 12 | 326 | 29 | **sTILs** | 3 | 16 | 321 | 30 |

Supplementary Table 4. Confusion matrices evaluated on the inferred P-truth testing dataset for models trained under different train/val splits, as well as the ensemble model.

|  | **Feature Names** | **Description** |
| --- | --- | --- |
| Clinical features | Age at diagnosis  AJCC tumor stage  ER status by IHC  PR status by IHC  HER2 status by IHC | Age at Diagnosis  AJCC pathologic tumor stage  Estrogen Receptor (ER) status by immunohistochemistry (IHC)  Progesterone Receptor (PR) status by immunohistochemical (IHC)  Human Epidermal growth factor Receptor 2 (HER2) by immunohistochemical (IHC) |
| TME features based on Delaunay graph | t.count.prob  s.count.prob  l.count.prob  t_t.edges.marginal.prob  t_s.edges.marginal.prob  t_l.edges.marginal.prob  s_s.edges.marginal.prob  s_l.edges.marginal.prob  l_l.edges.marginal.prob  t_s.edges.conditional.prob  t_l.edges.conditional.prob  s_t.edges.conditional.prob  s_l.edges.conditional.prob  l_t.edges.conditional.prob  l_s.edges.conditional.prob | Percentage of tumor nuclei  Percentage of stromal nuclei  Percentage of sTILs nuclei  Percentage of tumor-tumor connections over all nuclei connections  Percentage of tumor-stromal connections over all nuclei connections  Percentage of tumor-sTILs connections over all nuclei connections  Percentage of stromal-stromal connections over all nuclei connections  Percentage of stromal-sTILs connections over all nuclei connections  Percentage of sTILs-sTILs connections over all nuclei connections  Percentage of tumor-stromal connections over all stromal connections  Percentage of tumor-sTILs connections over all sTILs connections  Percentage of stromal-tumor connections over all tumor connections  Percentage of stromal-sTILs connections over all sTILs connections  Percentage of sTILs-tumor connections over all tumor connections  Percentage of sTILs-stromal connections over all stromal connections |
| TME features based on nuclei densities | t.norm  s.norm  l.norm  t_s.cos  t_l.cos  s_l.cos  t_s.proj.prob  s_t.proj.prob  t_l.proj.prob  l_t.proj.prob  s_l.proj.prob  l_s.proj.prob | Density of tumor nuclei in tissue  Density of stromal nuclei in tissue  Density of sTILs nuclei in tissue  Degree of tumor-stromal interaction intensity  Degree of tumor-sTILs interaction intensity  Degree of stromal-sTILs interaction intensity  Degree of stromal tissue invaded by tumor  Degree of tumor tissue surrounded by stromal  Degree of sTILs surrounded by tumor  Degree of tumor tissue invaded by sTILs  Degree of sTILs surrounded by stromal  Degree of stromal tissue surrounded by sTILs |

Supplementary Table 5. Features included in survival analysis and their meanings. Clinical features are defined in the original BRCA meta-information table. TME features based on Delaunay graphs in random patches of the ROI are defined in HD-Staining. TME features based on nuclei densities are defined by HD-Yolo in section 2.2.2.

|  | IDC only model | | | IDC + ILC model | | |
| --- | --- | --- | --- | --- | --- | --- |
| Selected features | p-value | importance | coefficients | p-value | importance | coefficients |
| Age at diagnosis | 3.566e-05 | 0.0961 ± 0.0920 | 4.5270 | 1.756e-05 | 0.0875 ± 0.0686 | 2.5552 |
| AJCC tumor stage | 2.254e-03 | 0.1048 ± 0.0820 | 0.5623 | 1.981e-03 | 0.1153 ± 0.0796 | 0.5206 |
| ER status by IHC | 4.430e-01 | -0.0150 ± 0.0292 | -0.1690 | 3.824e-01 | 0.0013 ± 0.0229 | -0.2172 |
| PR status by IHC | 2.005e-01 | 0.0071 ± 0.0370 | -0.2629 | 9.096e-02 | 0.0075 ± 0.0391 | -0.2855 |
| HER2 status by IHC | 6.927e-01 | 0.0011 ± 0.0054 | 0.0149 | 2.009e-01 | 0.0171 ± 0.0397 | 0.2210 |
| t_l.proj.prob | 9.640e-03 | 0.0249 ± 0.0130 | 0.3704 | 2.810e-03 | 0.0801 ± 0.0407 | 0.9735 |
| l_t.proj.prob | 4.202e-02 | 0.0088 ± 0.0166 | -2.8152 | 1.838e-02 | 0.0041 ± 0.0069 | -0.4691 |

Supplementary Table 6. P-values, feature importance, and coefficients for ElasticNet-selected features.
